# Protocol to optimize gain settings of Aurora for better data resolution

**DOI:** 10.1101/2025.06.11.659153

**Authors:** Debajit Bhowmick, Lindsay Richard, Michelle Leigh Ratliff

## Abstract

Full-spectrum flow cytometry has demonstrated clear advantages over conventional approach. Cytek’s Aurora platform has played a pivotal role in popularizing this technology, even though SONY introduced a full-spectrum flow cytometer earlier. Full spectral cytometers can identify the autofluorescence and unmix the autofluorescence much better than conventional cytometers. The default gain setting for Aurora, known as the Cytek Assay Setting (CAS), is effective for its intended purpose. However, there is no clear evidence to confirm that CAS provides optimal data resolution. This technical note outlines a methodology for enhancing gain settings to achieve a two-to four-fold improvement in resolution on average.

## Introduction

The major utility of any flow cytometer is to run Immunophenotypic panels. A critical step in any polychromatic flow cytometry experiment is reagent titration, which ensures optimal separation between negative and positive populations. This step is particularly significant for identifying cells with dim antigen expressions. Beyond reagent titration, it is equally essential to optimize detector sensitivity by maximizing the signal-to-noise ratio (SNR), as this further enhances resolution. Maximizing SNR will make sure dim positives are visible over the negative cells. Successful separation of the dim population will significantly impact the high dimensional data analysis outcome. With optimal SNR researchers will be able to identify population which are hard to distinguish, especially in exploratory work. Established protocols exist for detector sensitivity optimization in photomultiplier tube (PMT)-based systems (1,2) and avalanche photodiode (APD)-based systems such as CytoFlex (3).

Recent studies, including OMIP-95 (4), have proposed methods to optimize gain settings for the Aurora platform. However, OMIP-95 did not show any direct comparison between cell populations identified by CAS and their high-sensitivity settings. In this work, we present a simple procedure for optimizing Aurora’s gain settings to achieve superior data resolution and separation.

## Material and Method

To determine the optimal gain settings, we performed a thorough gaintration. The first step involves assessing the intrinsic noise levels of all fluorescence detectors across various gains. This was achieved by blocking the collection objective and optical fiber opening with two pieces of aluminum foil, as illustrated in Figure 1A. This setup allowed the detection of the forward scatter (FSC) signal while prevent any fluorescence reaching the detectors.

**Figure 1:**
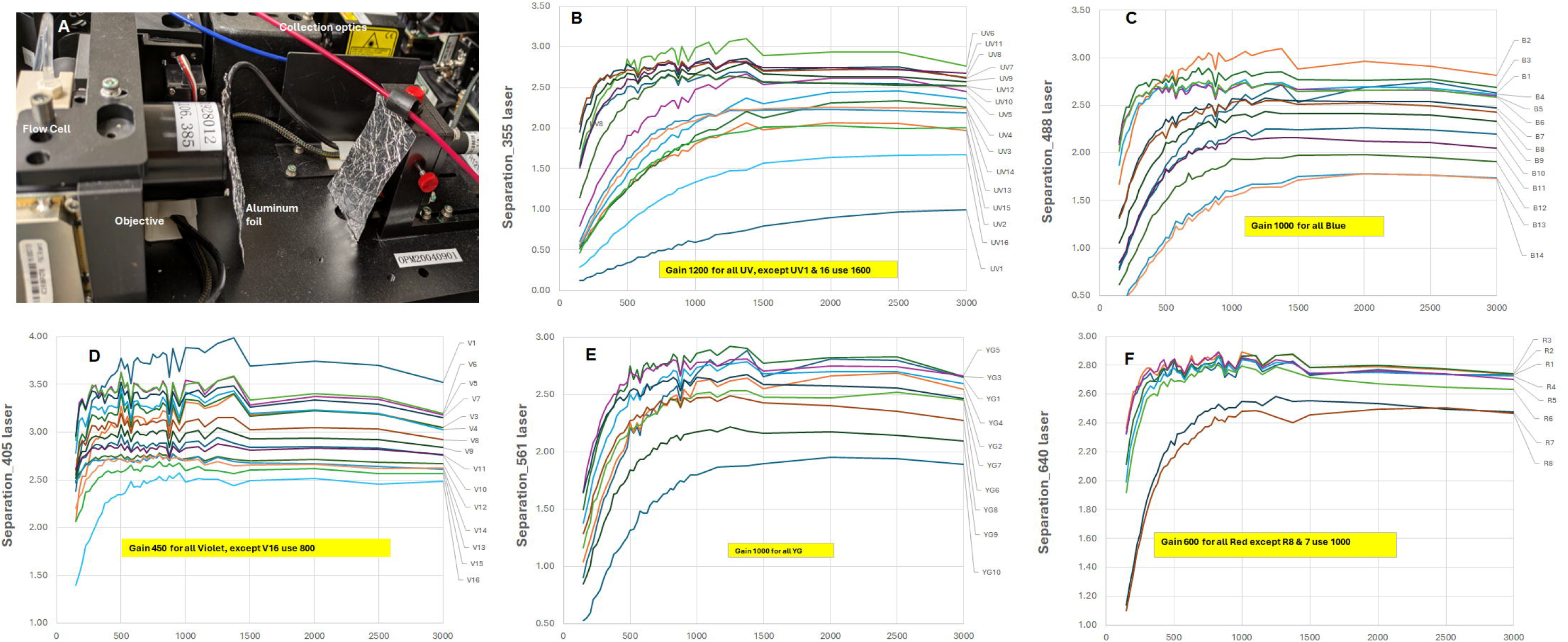
Light blocking strategy. (A) The placement of two pieces of aluminum foil was implemented at the openings of the collection objective and optical fibers to block fluorescence light from reaching the avalanche photodiode (APD) detectors. (B–F) These panels illustrate the separation values obtained for all detectors under the five laser configurations. The gain values highlighted in yellow represent the selected gain settings for corresponding detector. These gain values were chosen based on their position just below the plateau in the separation value curves, ensuring optimal performance without saturation.

For every fluorescence detector, gain values were adjusted manually to a predetermined level, such as 150. Using the FSC histogram, we recorded 10,000 peripheral blood mononuclear cells (PBMCs) from the main population. The procedure was then repeated for subsequent gain values, such as 175, 200 and continued incrementally up to 3000.

After collecting the noise data (blank), the same process was repeated without the aluminum foil to measure cellular autofluorescence (AF). This step allows other than detector’s noise to be collected. From the resulting .fcs files, the median fluorescence intensity (MFI) and robust standard deviation (rSD) were extracted for the PBMCs. The gaintration values (separation parameter) were quantified using the following formula.

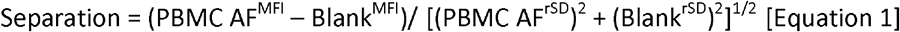

Figures 1B to 1F illustrate the patterns observed for all detectors of five lasers. From these graphs, a gain value was selected for each detector at a point just below the plateau in the respective curve. These selected values are indicated in Figure 1 and collectively constitute the high-sensitivity gain setting, referred to as HSen.

To evaluate the performance of HSen in a real life experiments, a 15-color panel was designed (details provided in Supplementary Table 1). Antibody titration was conducted using the CAS, including the Live/Dead amine-reactive dye. PBMCs from human donors were used to prepare all single stains, with unstained cells serving as the unmixing reference. Data acquisition, unmixing, and analysis were performed using SpectroFlo software (v2.2), without any manual adjustments to the unmixing matrix. AF extraction was not performed. Data was collected from eight separate donors over eight different days, and the dataset has been made publicly available on the figshare.com (https://doi.org/10.6084/m9.figshare.28528361.v1).

PBMCs from a young healthy male adult were isolated from a leukocyte reduction chamber (LRC) purchased from the Oklahoma Blood Institute, with Lymphoprep (StemCell Technologies). Oklahoma Blood Institute performed relevant informed consents for blood product use for research purposes. Because the chambers are a byproduct of platelet donation and do not cause additional risk to the donor, the ECU Institutional Review Board does not require board review of protocols. Protocols were approved by East Carolina University Institutional Biosafety Committee protocol #01–19. All the methods were carried out in accordance with relevant guidelines and regulations.

Cells were stained with various antibodies as outlined in the preceding section. Prior to antibody addition, 5 μl of Brilliant Stain Buffer (BD, Catalog No. 563794) and 5 μl of True-Stain Monocyte Blocker (BioLegend, Catalog No. 426101) were added to each tube and incubated for 15 minutes. The final staining volume was standardized to 200 μl. The cells were then incubated with an antibody master mix for 30 minutes, in ice followed by two washes with 3 ml of cold PBS. Subsequently, the cells were stained with a Live/Dead dye on ice for an additional 15 minutes, washed twice with 3 ml of cold FACS buffer, and finally resuspended in 250 μl of FACS buffer. The samples were maintained on ice throughout the procedure.

Data acquisition (single stain and full stain) was conducted separately using both the CAS and the HSen. Corresponding single stains were used to unmix the full stain. Unmixed data were manually gated, and the MFI as well as the rSD were calculated for each gated population. A comparative parameter (resolution change) was used to evaluate the changes in data resolution achieved with HSen relative to CAS across various populations.

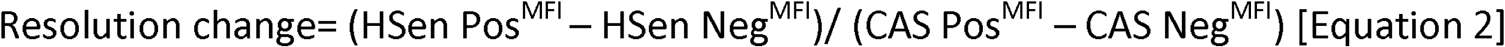

Apart from equation 2 we used Stain Index (SI) to calculate the change of the resolution.

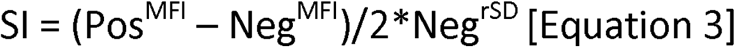

Increase of the rSD of the negative population due to HSen was calculated by

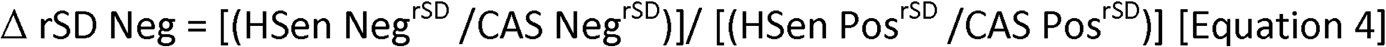

## Results

Figure 1 demonstrate that separation values increase sharply with increase of detector gain and reaches a plateau. For different detector the rate of rise is dissimilar, some are shallower than other. Based on the appearance of the plateau we choose the target gain to be slightly below of the plateau. Detectors with shallow rise has comparatively higher gain values from the rest of the detectors in the same array.

Supplementary Figure 1 provides examples demonstrating how data resolution improves when transitioning from the CAS to the HSen gain settings. Improvement in resolution between negative and positive populations are marked by red arrow while better resolution of the dim populations is marked by green arrow. Improved resolution for dim populations is evident by the rise of the frequency of the cells indicated by the heat map or by the appearance of the new populations. Due to logarithmic scaling, subtle resolution changes are not always visually apparent across all samples. To address this, a parameter (resolution change) was created to measure the improvement of the resolution, as depicted in Figure 2. The resolution change graph (blue line) highlights a 2-to 4-fold improvement for most fluorochromes with HSen, with CD14-BUV395 showing an 8-fold increase.

**Figure 2:**
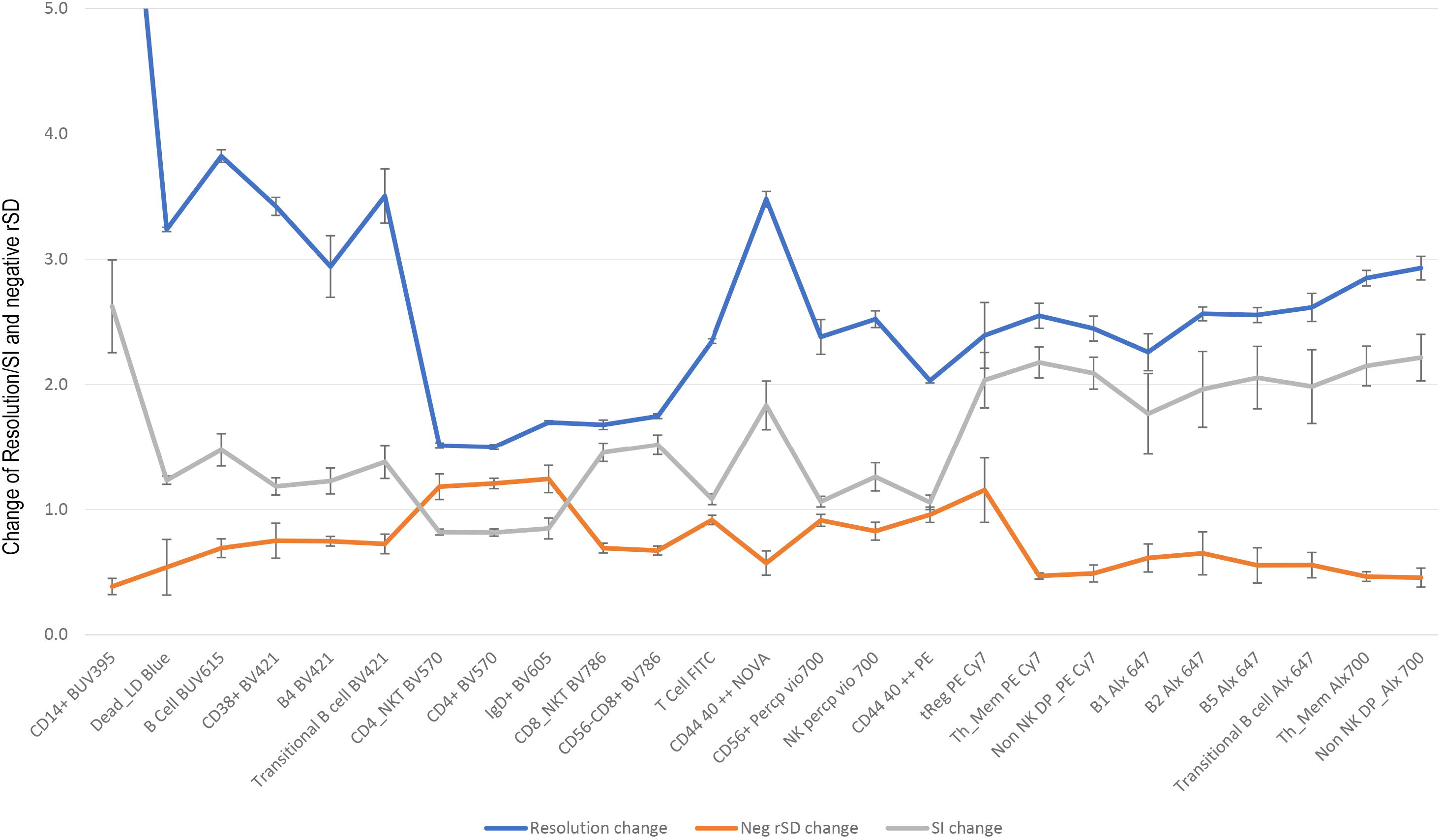
Effect of HSen on resolution and spread of negative population. The populations on the X-axis are organized based on the ascending wavelengths of the lasers, with populations within each laser further arranged according to their emission wavelengths. The blue line represents the change in resolution for each individual population. The gray line illustrates the Stain Index ratio between the HSen and CAS for the same populations. Additionally, the orange line depicts the relative increase in the robust standard deviation of the “negative” population.

Apart from the resolution change parameter we used the Stain Index parameter as well. We plotted SI^HSen^/SI ^CAS^ in Figure 2 (Grey line). Most of the markers showed increase in SI in HSen. Some marker exhibited minimal or no improvement (SI change ≤ 1) with HSen. SI increase on an average of 2 fold using HSen. Further investigation revealed that in these cases, the rSD of the negative population increased disproportionately under HSen. As we increased the gains for all detectors in HSen, it is natural that rSD itself will increase. This made it difficult to calculate whether the increase of the rSD in HSen is proportional to gain or not. To calculate the rSD rise we created a new parameter as per equation 4. The data was plotted in Figure 2 in Orange.

## Discussion

We initially utilized the Stain Index to identify the optimal SNR for each detector for gaintration. However, SI values continued to increase steeply up to a gain of 3000 without any sign of plateauing. It clearly indicates that only using the intrinsic detector noise (light blocking) is not enough to represent the noise of the cytometer. Upon further consideration, it became evident that the rSD of the cells AF need to be incorporated into the calculation. The AF signal does have all the other noise that is present in the cytometer. Inclusion of rSD revealed a plateau at certain gain values. Literature search showed James Wood had suggested same approach in his 1998 article (5). To ensure that signals will not be off scale we selected the gain values just under the plateau. In our previous paper (3) we showed that light blocking approach works with APD based CytoFlex manufactured by BeckMan Coulter.

Figure 2 shows negative cells has higher rSD for some markers using HSen. For CD4 BV570 and IgD BV605 the increased rSD showed drop in SI ratio and it also brings down the resolution change values as well. Non specific binding of the antibody and/or the fluorochrome itself introduced some very dim signal in the negative population, HSen setting picked up the signal. This observation indicates the HSen has much superior resolving power over CAS and can be useful to identify very dim signal over truly negative signal. We did run a titration using CAS setting. Optimizing titrations using HSen will address this issue and yield higher resolution.

During the gaintration we witnessed that SI is not ideal tool for this purpose. We are afraid that use of SI may bias our calculation with false positive manner. For that reason, we like to have another parameter to measure the change in data resolution. We created the resolution change parameter for this purpose. It identifies the ΔMFI for both settings and then calculates how much increase ΔMFI of HSen has over ΔMFI of CAS. The calculation does not include rSD component directly but as the Figure 2 showed MFI of the negative population will increase significantly with NSB and effect the resolution change values.

With both parameters we showed that HSen consistently exhibited higher resolution than CAS. We came up with the equation 4 to identify whether in HSen setting is picking up NSB in the negative population more than the CAS does. To do that we 1^st^ calculated the fold increase of the rSD of the negative in HSen over CAS. Then we calculate the same for positive population, because the panel is titrated, and samples were washed twice we can safely assume that there is no considerable NSB in the positive population. We divide these two to identify whether negative population have any extra rSD increase. This value is plotted in Orange in Figure 2. For CD4 BV570 and IgD BV605 we saw increase in the NSB to the negative population. It showed clearly that for CD4 and IgD markers has observable non specific binding in HSen setting. We have confirmed this in the corresponding 2D plots. Higer rSD rise of CD4 and IgD in HSen decreased the SI ratio under 1 and reduced the resolution change. Interestingly CD25 PE-Cy7 also showed rSD increase but both SI ratio and resolution change is higher than 1. This indicates the rSD increase does not necessarily bring down the separation of the population. SI is more sensitive to rSD increase than the resolution change parameter.

We believe HSen gain setting can be used to improve resolution for extracellular vesicles as well but that need to be tested. Folks who are not comfortable opening the machine can use non-fluorescent one-micron particles to get data similar to the blocked noise information we showed previously (3). These beads have almost no AF which means they can be used as an alternative of blank/noise information. Currently, no analytical tool exists to directly compare the effects of HSen and CAS settings on high-dimensional visualization and clustering. This limitation arises from the variability in MFI across fluorochromes, which complicates direct comparisons.

## Conclusion

HSen can provide superior data resolution compared to CAS. We do suggest performing reagent titration using HSen rather than CAS setting.

## Legends

**Supplementary Figure 1:**
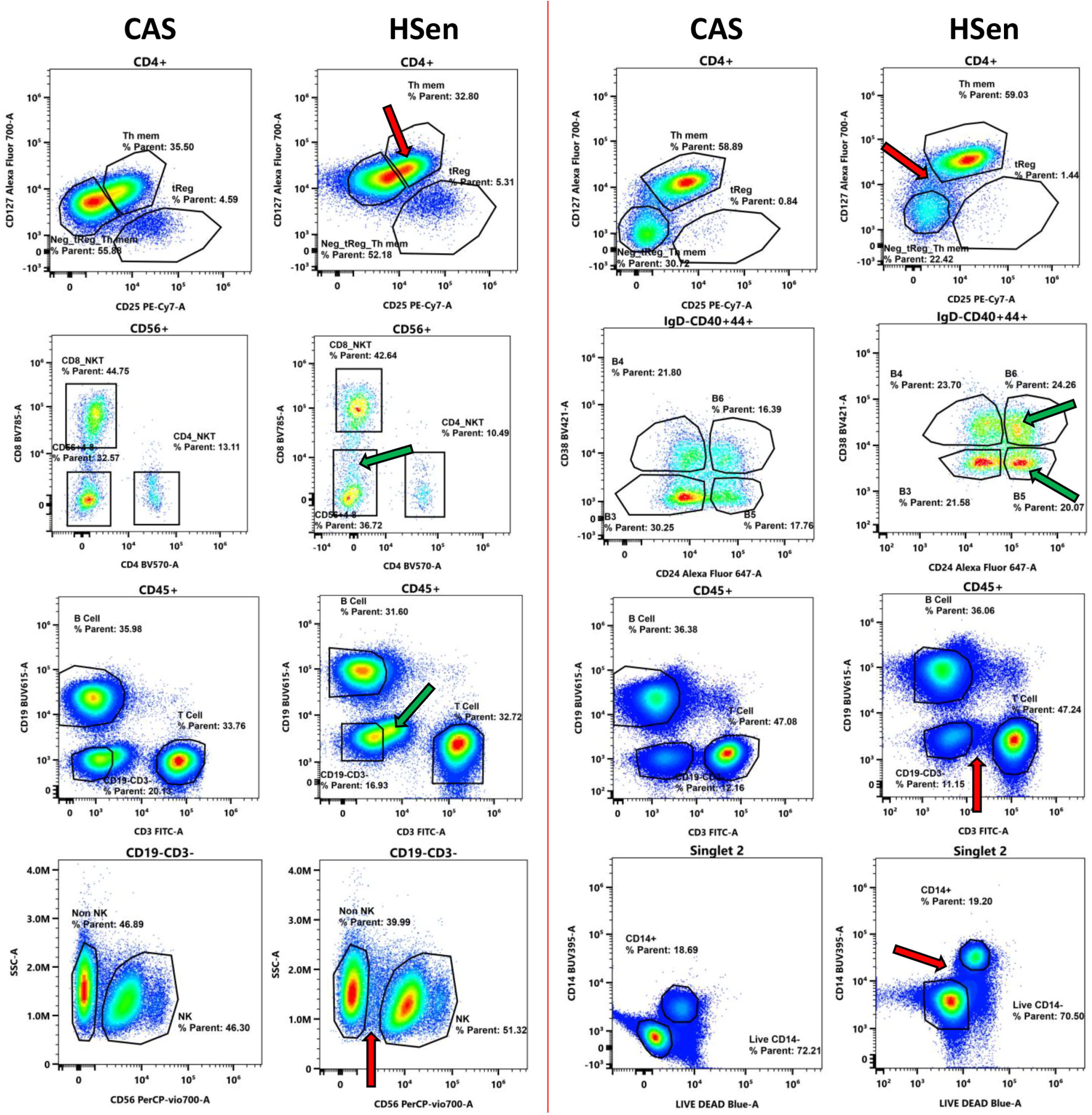
Examples of increased resolution. 2D plot distribution of the same sample ran using either CAS or HSen setting, showing higher separation on Hsen gain setting.

**Supplementary Table 1: Reagent details:** Catalog number, lot number and exact amount used of all the antibodies and Live Dead dye used in this study.

## Author contribution

DB and MLR contributed equally to write the text, conceptualize the original idea, performed data analysis. DB ran all samples. LR performed all the staining.

